# Muscle fibre size and myonuclear positioning in trained and aged humans

**DOI:** 10.1101/2023.08.25.551292

**Authors:** E. Battey, Y. Levy, R.D. Pollock, J.N. Pugh, G.L. Close, M. Kalakoutis, N.R. Lazarus, S.D.R. Harridge, J. Ochala, M.J. Stroud

## Abstract

Myonuclear domain (MND) is the theoretical volume of cytoplasm within which a myonucleus is responsible for transcribing DNA. Changes in myonuclear number, organisation, and myonuclear domain size are associated with exercise adaptations and ageing. However, data on satellite cell activation, changes in MND volumes and myonuclear arrangement following exercise are inconsistent. Additionally, whether MNDs and myonuclear arrangement are altered with age remains unclear. The aim of the present investigation was therefore to investigate relationships between age and activity status and myonuclear numbers and organisation. Muscle fibres from younger trained (YT) and older trained (OT) individuals were compared with age-matched untrained counterparts (YU and OU). Serial, optical z-slices were acquired throughout isolated muscle fibres and analysed to give 3D coordinates for myonuclei and muscle fibre dimensions, respectively. As expected, mean cross-sectional area (CSA) (μm^2^) of muscle fibres from OU was 29-42% smaller compared to the other groups. Number of nuclei relative to fibre CSA was 87% greater in OU compared to YU muscle fibres (P < 0.05). Additionally, scaling of myonuclear domain volume with fibre size was altered in older untrained individuals. Myonuclear arrangement, on the other hand, was similar across groups. These data indicate that regular endurance exercise throughout the lifespan may preserve the size of single muscle fibres in older age and maintain the relationship between fibre size and MND volumes. Inactivity, however, may result in reduced muscle fibre size and disrupted relationship between fibre size and MND volumes.

**Plain Language Summary:** In this study, we examined the relationship between physical activity and the characteristics of muscle fibres in individuals of different age groups. We focused on a concept called the myonuclear domain (MND), which refers to the volume surrounding muscle nuclei or myonuclei that house the genome. We wanted to understand how changes in myonuclear number, organisation, and MND size were influenced by exercise and aging. To do this, we compared muscle fibres from younger trained individuals, older trained individuals, and age-matched untrained individuals. The results showed that the average size of muscle fibres in the untrained older individuals was smaller compared to the other groups. Moreover, the number of nuclei relative to fibre size was significantly higher in the untrained older individuals. However, myonuclear arrangement was similar across all groups. These findings suggest that regular endurance exercise throughout life may help maintain muscle fibre size, myonuclear numbers, MND volumes, and myonuclear organisation in older individuals. Conversely, inactivity can lead to reduced muscle fibre size and disrupted relationship between fibre size and MND volumes. These results have important implications for understanding the effects of exercise and aging on muscle health.

## Introduction

Myonuclear domain (MND) volume is the theoretical volume of cytoplasm which a myonucleus is responsible for (Allen et al., 1999). Changes in myonuclear structure, number, organisation, and MND volume have been associated with muscle adaptation, including in response to exercise training, injury, disuse, disease, and ageing (Bagley et al., 2023; Battey et al., 2020, 2023; Mohiuddin et al., 2019; Murach et al., 2020; Stroud et al., 2017). However, data on changes in MND volumes and myonuclear arrangement following exercise are inconsistent (Abreu et al., 2017; Verney et al., 2008). Additionally, there are conflicting reports regarding the relationship between MNDs and myonuclear arrangement with ageing (Cristea et al., 2010; Karlsen et al., 2015).

One possible explanation for the inconsistency of findings related to changes in MND sizes, myonuclear number, and myonuclear arrangement are the use of transverse cross-sections and two-dimensional muscle fibre analysis. Specifically, transverse cross-sections of muscle biopsies from sedentary young, trained young, sedentary elderly and trained elderly individuals showed no differences in myonuclear numbers or domains between groups (Karlsen et al., 2015). Conversely, three-dimensional analysis of single muscle fibres revealed an age-related decline in MND volume, reduced fibre size, variability of MND volume and myonuclear spacing (Cristea et al., 2010). However, physical activity levels were not accounted for in this study, preventing delineation between effects of inactivity and ageing. Thus, greater clarity on the relationship between myonuclear number and positioning, ageing, and exercise could be achieved by i) analysing longitudinal segments of muscle fibres in 3D, and ii) considering the influence of activity levels on age-related observations.

We aimed to explore relationships between age and activity status and myonuclear numbers and organisation by analysing muscle fibres from younger trained (YT), older trained (OT), and untrained counterparts (YU and OU). Muscle fibres were stained with DAPI and phalloidin, followed by acquisition of serial optical z-slices through the fibre and quantification of 3D coordinates for myonuclei and muscle fibre dimensions, respectively. Here, we found that muscle fibres from OU contained more myonuclei relative to fibre size and a disrupted relationship between MNDs and fibre size. Additionally, smaller fibres from OU had relatively higher nuclear numbers and smaller MNDs compared to the other groups. No significant association between ageing and myonuclear arrangement was observed.

## Methods

Human participant characteristics and ethical approval Four mixed gender groups were recruited to participate in the current study (n = 6 per group). These groups were: younger untrained healthy (YU) (33 ± 9.5 years), younger trained marathon runners (YT) (32 ± 5.4 years), older untrained individuals (OU) (79 ± 11.3 years), and older highly trained master cyclists (OT) (75.5 ± 3.2 years) (table 2.1). Due to sample availability, marathon runners rather than cyclists were used for the YT group.

**Table 1.**
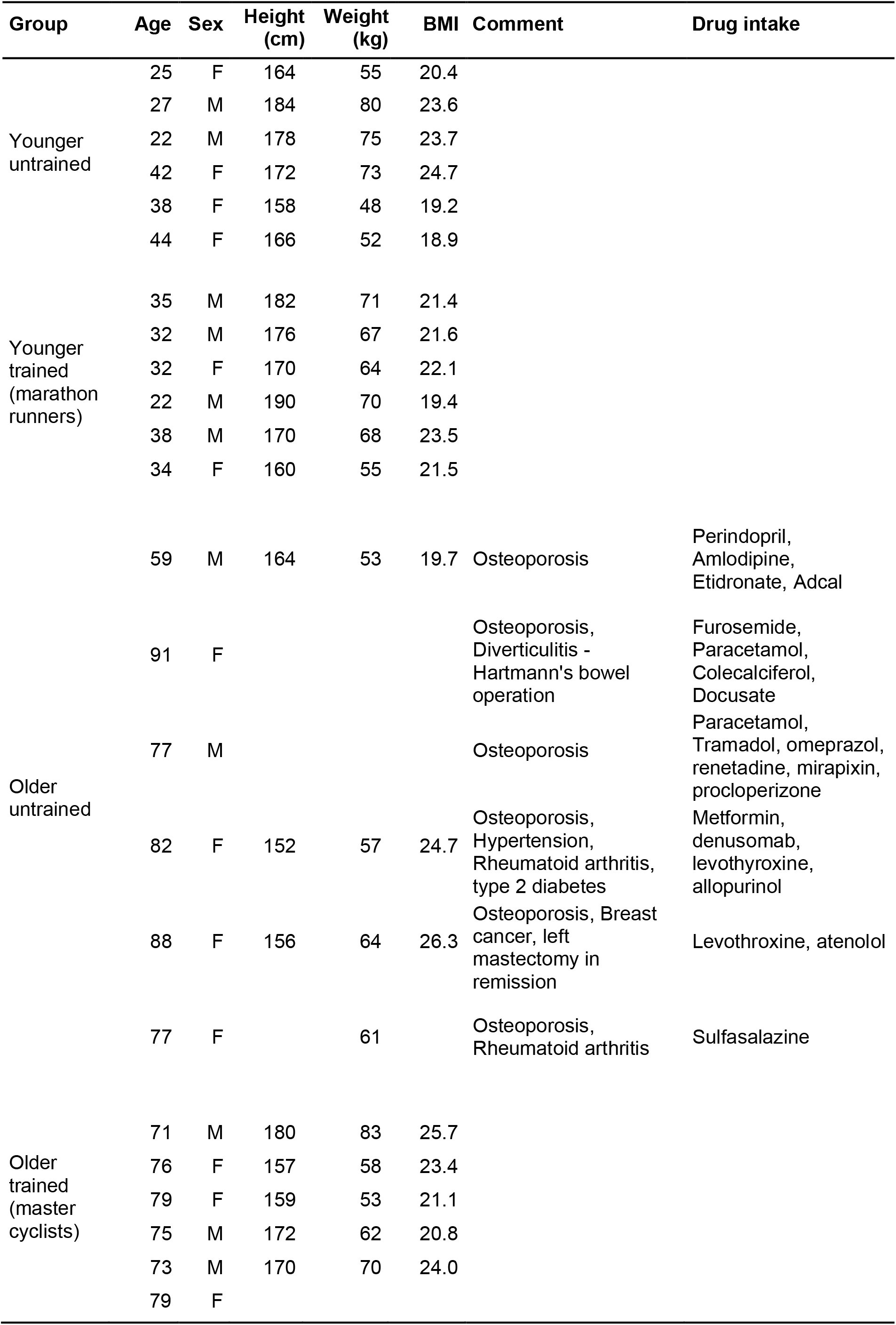
Participant characteristics.

The YU group was considered healthy, but not necessarily sedentary, as two of the participants had been participating in low-level recreational sport activities (<2 sessions/week) at the time of the study. Thus, the young cohort consisted of sedentary and low-level physically active individuals. Participants were considered healthy if they met the criteria outlined by (Greig et al., 1994). The YT group consisted of trained recreational marathon runners (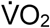 peak 56.7 ± 6.6 ml.kg.min^-1^, mean ± SD). In the YT group, the mean fastest running times (min, mean ± SD) in the previous 18 months over marathon, half marathon and 5 km were 204.5 ± 14.2, 88.5 ± 3.3, and 19.8 ± 1.3, respectively.

The OU group, used as a model for muscle disuse in old age, was a previously characterised cohort, who underwent dynamic hip screw insertion surgery (Kalakoutis et al., 2023). The patients completed a basic physical health questionnaire and were considered eligible if they did not suffer from neuromuscular disease. The hip fracture patients were recruited on the basis that they were likely to be frail individuals, and some had underlying health conditions (Table 1).

The OT group consisted of previously characterised individuals (Pollock et al., 2015, 2018) who were amateur master cyclists (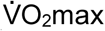 36.76 ± 8.56 ml.kg.min-1, mean ± SD). Master cyclists were included if they were able to cycle 100 km in under 6.5 hours (males) or 60 km in under 5.5 hours (females). Participants must have had completed this distance within the specified time on two occasions in the three weeks prior to the date of participation in the study.

Prior to participation, written informed consent was obtained from all subjects. Procedures were approved by the Fulham Research Ethics Committee in London (12/LO/0457), Westminster Ethics Committee in London (12/LO/0457) or Liverpool John Moores ethics committee (H17SPS012) and conformed to the standards set by the Declaration of Helsinki. All human tissues were collected, stored, and analysed in accordance with the Human Tissue Act (World Medical Association, 2013).

### Obtaining muscle samples and isolating single muscle fibres

*Vastus lateralis* muscle samples from the YU, YT, and OT participants were obtained using the same biopsy procedure. A portion of the mid-thigh was shaved, and the skin was cleaned using chlorhexidine gluconate. 2% lidocaine was applied to the skin prior to making a 5 mm incision in the skin using a scalpel. A Bergström biopsy needle was inserted into the incision and a biopsy of approximately 200 mg was taken. Approximately 60 mg of the biopsy sample was then placed in relaxing solution (Table 2) in a petri dish on ice. Muscle tissue from the OU were obtained during dynamic hip screw insertion surgery. During the exposure of the hip during surgery, a portion of the *vastus lateralis* that becomes detached was placed into ice-cold relaxing solution. Muscle samples from biopsies and hip fracture surgeries were then prepared for skinned fibre experiments.

Following excision, muscle samples (submerged in relaxing solution in a petri dish) were divided into bundles of approximately 100 muscle fibres using forceps under a stereo microscope (Zeiss, Stemi 2000-C) with a separate light source (Zeiss Stereo CL 1500 ECO). The ends of the bundles were then tied onto glass capillary tubes using surgical silk (LOOK SP102) and stretched to approximately 110% of the original length. These bundles were subsequently placed into 1.5 ml Eppendorf tubes, containing skinning solution (relaxing solution with 50% (v/v) glycerol), at 4°C for 24 h to permeabilise the muscle fibres by disrupting the lipid bilayer of the sarcolemma, leaving myofilaments, intermediate filaments, and nuclear envelope intact (Frontera & Larsson, 1997; Konigsberg et al., 1975; Stienen, 2000; Wood et al., 1975). Samples were then treated in ascending gradients of sucrose dissolved in relaxing solution (0.5 M, 1 M, 1.5 M, 2 M) for 30 min to prevent cryodamage (Frontera & Larsson, 1997). In a petri dish containing 2 M sucrose, fibres were then removed from the glass capillary tubes before being placed in cryovials and snap-frozen in liquid nitrogen.

For immunofluorescence experiments, muscle fibre bundles were placed in descending levels of sucrose dissolved in relaxing solution, for 30 minutes in each solution (2 M, 1.5 M, 1 M, 0.5 M, 0 M). Samples were transferred to skinning solution at -20°C until the day of an experiment. To normalise muscle fibre tension and orientation, single fibres were isolated from bundles using forceps and mounted on half-split grid for transmission electron microscopy (TEM) glued to a coverslip. This was process was repeated for each fibre so that an array of 9-11 fibres were mounted on the same platform.

### Immunostaining, imaging, and analysis of single muscle fibres

Muscle fibres were permeabilised in 0.2% triton for 10 min and fixed in 4% PFA for 15 min. Muscle fibres were then incubated in a solution containing DAPI (Molecular Proves, D3571; 1:800) and Alexa Fluor™ 594 Phalloidin (Invitrogen, A12381; 1:100) for one hour, to visualise myonuclei and Actin, respectively. Images were then acquired using a Zeiss Axiovert 200 confocal microscope equipped with a 20× objective and CARV II confocal imager (BD Bioscience, San Jose, CA). 100 images were acquired, in 1 μm z-increments, centred around approximately the middle of the fibre to ensure the entire depth of the fibre was imaged. Finally, mounting medium (Fluoromount-G^®^) was added to the coverslip and another 22 × 50 mm coverslip was placed carefully (to avoid bubbles) on top. Between each step of staining protocols, fibres were washed four times in PBS, leaving the fibres submersed in the final PBS wash for 5 min.

Three-dimensional visualisation of muscle fibres and quantification of myonuclear arrangement was carried out using Metamorph software. Z-stacks were imported, and each nucleus within the field of view (using DAPI signal) and muscle fibre edges in XY and Z planes (using phalloidin signal) were recorded on Excel (linked to Metamorph). These data were analysed using a MATLAB script to obtain MND (volume of cytoplasm governed by each nucleus), order score (spatial distribution of nuclei), nearest neighbour distance (distance between nuclei) and fibre dimension values (Bruusgaard et al., 2003; Levy et al., 2019). Order score was calculated by generating a theoretical optimum distribution (MO) and a theoretical random distribution (MR) for each muscle fibre, based on the coordinates and numbers of corresponding myonuclei. This was compared with the experimental distribution (ME) and an order score was calculated with the equation: (ME−MR)/(MO−MR).

### Statistical analyses

To analyse whether an overall significant difference was present between muscle fibres from the different human groups, two-way analysis of variance (ANOVA) tests were carried out, followed by post-hoc Tukey tests to specify between which groups differences existed. Mean values for each individual were used for two-way ANOVA. For correlation analyses, a simple linear regression was performed to test if slopes were significantly non-zero and nonlinear straight-line regression analyses were performed to compare the slopes of different conditions. For all statistical tests, P < 0.05 indicated significance (* P < 0.05, ** P < 0.01, *** P < 0.001). Error bars in graphs represent mean ± SD. All data were statistically analysed using Prism 9 (GraphPad).

## Results

### Scaling of myonuclear domain volume with fibre size is altered in older untrained individuals

To investigate the effects of ageing and exercise on myonuclear organisation and number, myonuclear domain (MND) volumes, myonuclear arrangement, and fibre dimensions were compared in muscle fibres from YU, YT, OU, and OT (136, 83, 96, and 88 fibres analysed per group, respectively; 403 fibres analysed in total). As expected, due to the known effects of inactive ageing on muscle mass (Mckendry et al., 2018), the mean cross-sectional area (CSA) (μm^2^) of muscle fibres from OU was 29-42% smaller compared to the other groups (OU, 1980 ± 1105; YU, 4238 ± 1105; YT, 2941 ± 594; OT 2985 ± 626; YU vs OU P <0.05; Figure 1, 2A). Although the number of nuclei per fibre length (count/ mm) was unaffected by age or training status, number of nuclei relative to fibre CSA was 87% greater in OU compared to YU muscle fibres (24.1 ± 9.6 vs 12.7 ± 2.9, respectively, P < 0.05; Figure 2C).

**Figure 1.**
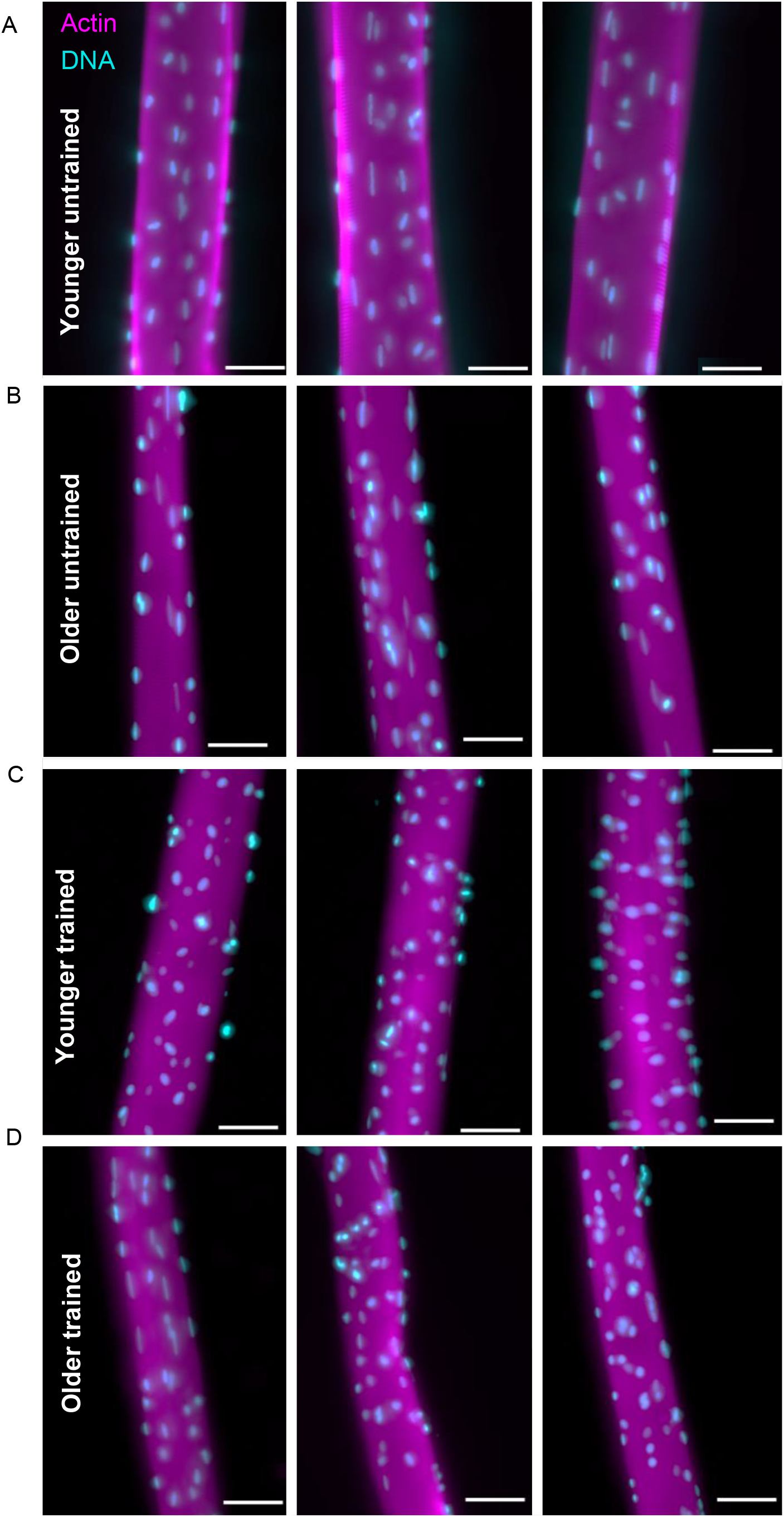
Representative images of myonuclear organisation in younger and older untrained and endurance-trained individuals. Representative images of vastus lateralis muscle fibres isolated from younger untrained (A), older untrained (B), younger trained (C) and older trained (D) stained with DAPI (cyan) and Phalloidin (magenta) antibody to visualise myonuclei and Actin, respectively. Scale bars, 50 μm.

**Figure 2.**
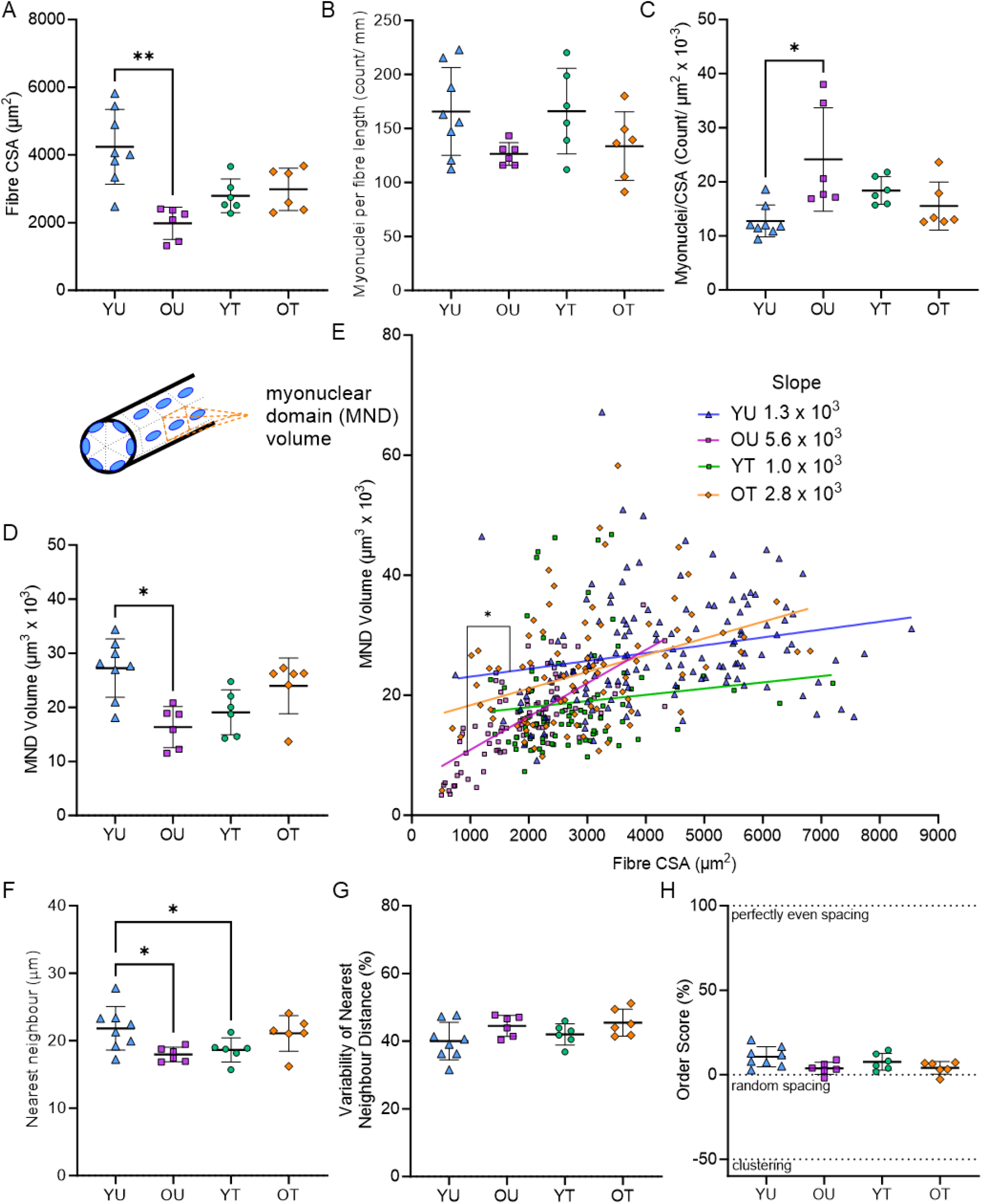
Muscle fibre size and myonuclear number, domains, and organisation in younger and older untrained and endurance-trained individuals. (A-C) Comparisons of muscle fibre cross-sectional area (CSA, μm^2^), myonuclei per fibre length, and myonuclei normalised to fibre cross-sectional area in muscle fibres from younger untrained (YU), older untrained (OU), younger trained (YT) and older trained (OT) individuals (D) Comparison of mean MND volumes across groups. Inset shows cartoon of MND volume, with orange dotted lines highlighting the MND of a single myonucleus (E) Relationship between muscle fibre CSA and myonuclear domain (MND) volume (μm^3^ x 103) in YU, OU, YT and OT. Coloured symbols represent individual muscle fibres. Simple linear regression and straight-line non-linear regression performed to quantify and compare gradients of slopes, * indicates P < 0.05 (F-G) Comparison of nearest neighbour distances (μm) and variability of nearest neighbour distances (%) across groups (H) Comparison of mean order scores (%) across groups. Dotted lines at -50, 0, and 100% represent clustered, random, and perfectly even spacing of myonuclei. (A-D, F-H) Two-way ANOVA used to compare mean differences between groups. Error bars represent mean ± SD.

In line with this, analysis of the volume of cytoplasm each myonucleus occupies (MND; μm^3^) revealed smaller mean MNDs in OU compared to YU (16.3 ± 3.8 vs 27.2 ± 5.4, respectively, P < 0.05; Figure 2D-E). Myonuclear domains were smaller in YT compared to YU (19.1 ± 4.14 vs 27.2 ± 5.4, respectively, P < 0.05). Further analysis of the relationship between fibre CSA (μm^2^) and MND volumes indicated that as fibre size increased, myonuclear domain volume (μm^3^) became larger in muscle fibres from OU compared to YU (Figure 2E; R^2^: YU, 0.05; OU, 0.47; YT, 0.01; OT, 0.15).

In addition to myonuclear number and MNDs, myonuclear arrangement was quantified. Consistent with smaller MND volumes, the mean distance between myonuclei (nearest neighbour, μm) was reduced in OU compared to YU, and in YT compared to YU (Figure 2F, P < 0.05). A previous report suggested that variability of nearest neighbour distance increased with age (Cristea et al., 2010). However, variability of nearest neighbour distance (%) was comparable between groups (Figure 2G). To gain further insight into the spatial arrangement of myonuclei, an order score (%) was generated to compute the distance between nuclei and fibre dimensions in X, Y and Z planes (Bruusgaard et al., 2003; Levy et al., 2018). An order score of 100% indicates completely even spacing, 0% random arrangement, and negative scores clustering. Order scores were similar across all groups (YU: 10.7 ± 6.0; YT: 10.1 ± 5.3; OU: 3.8 ± 3.6; OT: 4.1 ± 3.8) indicating comparable myonuclear arrangement (Figure 2H).

Together, these data show that muscle fibres from OU are smaller, have more nuclei per fibre CSA (μm^2^) and a greater expansion of the myonuclear domain as fibre size increases compared to muscle fibres from YU, YT, and OT. Myonuclear arrangement, on the other hand, was similar across groups.

## Discussion

A major problem in human ageing research and in skeletal muscle biology is to separate the effects of ageing from the negative effects of lifestyle factors that impact the ageing process (Distefano & Goodpaster, 2018; Lazarus & Harridge, 2017; Miljkovic et al., 2015) with one of the most documented being a lack of physical activity/exercise. It is widely recognised that the default position for humans is to be physical activity and/or exercise (Blair et al., 1989; Booth et al., 2012; Gries et al., 2018; Lazarus & Harridge, 2017; Pedersen & Saltin, 2015). In the absence of exercise, the ageing trajectory intersects with disease in and around the fifth to sixth decade of life. Thus, without exercise underlying physiological processes are essentially pathological while with exercise the ageing process, although in continual decline, is dependent upon the maintenance of physiological processes.

The aim of the present study was to determine the association between age and activity status regarding the number and organisation or nuclei within single human skeletal muscle fibres. We did this by studying fibres obtained from four groups individuals representing a spectrum of age and exercise status. The main findings were that in OU, MNDs were smaller and myonuclear number relative to fibre size was greater than YU. Additionally, MND volume in OU expanded with increasing fibre size to a greater extent than YU. Furthermore, we did not observe an association between ageing and myonuclear arrangement.

The higher number of myonuclei in smaller fibres in OU indicate that myonuclei were retained during age-associated disuse atrophy. If net loss of myonuclei contributed to muscle atrophy, one would expect myonuclear number to decrease with fibre size. However, we observed that myonuclear number was maintained in smaller fibres, which is consistent with the observation of others (Malatesta et al., 2009; Snijders et al., 2020). These data support the concept of ‘muscle memory’, whereby myonuclei are not lost with age or disuse atrophy (Gundersen, 2016; Gundersen et al., 2018). However, recent evidence demonstrating myonuclear loss and turnover suggests that new myonuclei could have been added to OU muscle fibres to maintain nuclear numbers (Kirby & Dupont-Versteegden, 2022; Murach et al., 2020; Snijders et al., 2020). Loss of myonuclei has only been demonstrated in mice (Murach et al., 2020; Snijders et al., 2020), so it is possible that this does not occur in humans, and that the myonuclei in OU fibres have been retained throughout the lifespan. However, we cannot rule out the possibility of this occurring in humans and future longitudinal studies of disuse are needed to test this hypothesis.

Our data highlight that myonuclear number alone is inadequate to maintain fibre size and suggest that other factors such as exercise-mediated gene expression and epigenetic changes may be required to maintain muscle fibre size and quality (Sharples et al., 2018; Widmann et al., 2019).

We observed that endurance-trained individuals did not have higher myonuclear numbers compared to untrained individuals, suggesting myonuclear accretion may not be a major endurance training adaptation. This is in line with previous evidence showing no increase in myonuclear number with endurance training in humans (Charifi et al., 2003; Snijders et al., 2011; Verney et al., 2008), despite reports of increased satellite cell activation and number after endurance training (Appell et al., 1988; Charifi et al., 2003; Verney et al., 2008). These results also demonstrate that lifelong aerobic cycling exercise in older age does not affect the number of myonuclei in single muscle fibres, supporting the results of a previous study in trained runners (Karlsen et al., 2015). Interestingly, we observed reduced MND volume in YT compared to YU, but this may be explained by the ∼25% smaller mean fibre size (not significant) resulting in a relative decrease in nuclear spacing.

Collectively, these data suggest that unlike the increase observed after resistance training (Petrella et al., 2008), endurance training is not associated with a substantial increase in myonuclear number. Thus, it is tempting to speculate that with endurance training, myonuclei adapt their epigenetic and transcriptional landscapes versus nuclear accretion as observed in resistance-trained fibres.

No changes in myonuclear organisation were observed with age, contrasting a previous report that showed higher variability of myonuclear positioning in type I fibres of older participants (Cristea et al., 2010). In the present investigation, some clustering and myonuclear disorganisation was observed in all groups, but these had less overall weight on the average values obtained. Thus, it appears that myonuclear organisation may be altered in some fibres in older groups, but not significantly more than other groups. A limitation of the present study is that in the interest of increasing the throughput of staining and imaging to acquire data on sufficient muscle fibres for statistical analyses, a fibre type co-staining was not performed. This additional fibre-type specific analysis would be required to directly address whether differences in fibre type proportions across groups affected myonuclear numbers and arrangement.

We observed no difference in mean fibre CSA in the older trained group compared to the other groups, in contrast to a recent study on lifelong endurance exercisers (Grosicki et al., 2021). This may be explained by the pooling of fibre types, with lifelong endurance exercisers from this cohort having a high distribution of smaller type I fibres (Pollock et al. 2018). Additionally, the YU group studied may have had a greater proportion of type IIA fibres compared to the OT group, as recently shown in Kalakoutis et al., (2023). Another possible explanation for the greater fibre CSA of some individuals in the YU group compared to the trained groups is that smaller fibres in endurance-trained individuals could be an adaptive response; efficiency is key for endurance performance and added muscle mass might increase the weight carried at the expense of efficiency (Coggan et al., 1992; Hughes et al., 2018; Joyner & Coyle, 2008). Additionally, increased muscle fibre size may increase diffusion distance between capillaries and mitochondria. This could increase the time it takes for oxygen to reach mitochondria, which may limit the rate of ATP production through aerobic metabolism, thereby compromising endurance performance. Lifelong endurance training can attenuate age-related muscle atrophy, which could explain the lack of significant difference in fibre CSA between YU and OT and significantly smaller CSA of muscle fibres from OU compared to YU (Wroblewski et al., 2011; Mckendry et al. 2018).

Our results on nuclear number and organisation also need to be interpreted in the context of our previous findings on these fibres regarding the shape of individual nuclei (Battey et al. 2023). Here we observed that fibres from both young and old untrained individuals were less spherical, and nuclei from older untrained were more deformable, and contained a thinner nuclear lamina than those from the older trained individuals. However, in contrast to the known adaptability of inactive young muscle fibres to exercise training, adaptation to exercise is more limited in older muscle, for example, anabolic resistance to exercise (Kumar et al., 2009) and feeding (Cuthbertson et al., 2005). This suggests that despite exhibiting some similar nuclear organisation features to YU, YT and OT, fibres from OU are compromised by an interaction of ageing and disuse, such as altered nuclear shape and mechanics (Battey et al., 2023).

We acknowledge that whilst we attempted to create a model in which age and exercise interactions could be studied, there are some limitations given the donors from which fibres were collected. These include: i) relatively small sample sizes that were not equally balanced between groups for sex; ii) whilst endurance exercisers were studied, in the YT group these were runners whereas they were cyclists in the OT, iii) the OU group were not simply healthy older individuals who were less active than OT, but older patients undergoing hip surgery who were also on different medications (Table 1), with the effects of this on muscle not fully understood and iv) the YU group, whilst being untrained had varying levels of physical activity, perhaps in part explaining the greater heterogeneity of fibre size in this group.

With these caveats in mind, the data do however, suggest that regular endurance exercise throughout the lifespan preserves the size of single muscle fibres in older age and maintains the relationship between fibre CSA and MND volumes seen in young, trained individuals. Young individuals who remain inactive are likely to be on a trajectory towards longer term pathology which may include reduced muscle fibre size despite maintained myonuclear number. Furthermore, in addition to myonuclear number and organisation, myonuclear structure and mechanics (Battey et al., 2023) are also likely to contribute to both reduced muscle fibre size and other transcriptionally regulated aspects of muscle associated with inactive ageing. Ideally, longitudinal studies comparing untrained and trained individuals who are more tightly defined will shed greater light on the interaction between ageing and inactivity processes in muscle.

## Additional Information

### Author Contributions

MS and JO contributed to the conception of the work. EB, JO, YL designed experiments. EB, YL, and JO performed experiments. EB and YL completed the formal analysis of the data. EB interpreted the data and completed data visualisation. RDP, MK, JNP, GLC, NRL, SDRH and JO recruited human participants and collected human muscle biopsy samples. EB wrote the first draft of the manuscript. All authors approved the final version of the manuscript to be published and agreed on all aspects of the work.

